# Strong segregation promotes self-destructive cooperation

**DOI:** 10.1101/2024.10.15.618393

**Authors:** Lingling Wen, Yang Bai, Yunquan Lan, Yaxin Shen, Xiaoyi She, Peng Dong, Teng Wang, Xiongfei Fu, Shuqiang Huang

**Author notes:** L. W. and Y. B. contributed equally to this work. Author contributions: X. F., and S. H. conceived and designed the experiments; L. W., Y. S., and X. S. performed the experiments; L. W., Y. B., P. D., and X. F. developed the models; L. W., and Y. B. analyzed the data; L. W., and Y. L. contributed materials/analysis tools; L. W., Y. B., P. D., T. W., X. F., and S. H. wrote the paper.

## Abstract

Self-destructive cooperator (SDC), where individuals sacrifice themselves for others, poses significant personal costs yet is widespread in nature. Traditional group selection theory struggles to explain its persistence, as the extreme cost of self-destruction typically exceeds the benefit to these individuals. Here, we predict that this behavior can endure in structured environments with strong segregation, where populations are divided into homogenous groups originating from one or two founding members. In such contexts, the benefits derived from SDC are primarily confined within the homogenous SDC groups, thus preserving the value of this altruistic sacrifice and ensuring its maintenance. To validate our hypothesis, we employ a synthetic system with engineered bacteria to mimic both SDC and non-sacrificing cheaters. We then conduct automated experiments facilitated by biofoundry technology to monitor and operate the subgroups with diverse growth behaviors stemming from a restricted number of initial cells because of strong segregation. Eventually, we demonstrate that SDC indeed maintains under strong segregation, and high stress benefits SDC because it reduces the benefits received by cheaters in heterogenous subgroups. This study extends the group selection theory to encompass even the most extreme manifestations of altruism and highlights the potential of automation in evolutionary research.

## Introduction

Self-destructive cooperation represented a high-level manifestation of altruism, where an individual sacrificed himself to produce public goods that ultimately benefited all other members of society^1-5^. Such noble behavior was not limited to humans with heightened consciousness and was also evident in more primitive organisms^6^. Examples included *Escherichia coli* expelling colicins to protect kin^7^, honeybees stinging to defend their hives^8^, and the immune system’s sacrificial response to sepsis^9^. Such self-destructive cooperative behavior in primitive organisms posed a profound evolutionary puzzle: Since self-destructive cooperation inherently reduced individual fitness, how could they persist in natural selection, a process driven by the imperatives of survival and reproduction.

Traditional group selection theory had successfully explained the maintenance of mild cooperative behaviors of primitive organisms under structured environments. This was predicated on the notion of “multi-level selection”, that cooperators could enhance the productivity of subgroups with a higher composition of cooperators, yielding a net advantage at the group level despite the intra-group disadvantage compared to non-cooperating counterparts^10-13^. However, such a theory failed to explain the evolution of extreme cooperation such as self-destructive cooperation, given that the self-destructive cooperators (SDCs) completely sacrificed themselves, leaving no offspring in subgroups, creating an extreme disadvantage within the group, where potential benefits at the group level might not compensate the cost^13^. Current theories suggested that self-destructive cooperation was not a directly selected trait but an unintended consequence of another beneficial function^14,15^.

In this study, we considered the evolution of self-destructive cooperation in a structured environment with ‘Segregation-Growth-Stress-Pool’ (SGSP) procedures, similar to a life cycle of biofilms where dense bacterial communities emerged from a limited number of initial cells and expanded until they confronted environmental stress^16-25^. In this scenario, the mixed population of SDCs and cheaters was partitioned into distinct subgroups and then cultivated until external stress was introduced to induce the self-sacrifice of SDCs that benefited the other individuals. Our theoretical analysis and numerical simulations showed that the ratio of SDC increased under strong segregation where group sizes were as minimal as approximately 1 or 2 individuals per subgroup. Notably, we found that the lower initial composition of SDC necessitated weaker segregation strength for its maintenance, and higher stress levels favored the persistence and proliferation of self-destructive cooperation.

These results were further demonstrated experimentally by employing synthetic SDC strains that offered a controllable model of self-destructive cooperation. Using automated experimentation from biofoundry technology, we overcame the challenge presented by the highly diverse growth dynamics inherent to tightly segregated subpopulations^26^. These findings not only extended classic group selection theory but also demonstrated the significant potential of biofoundries in advancing our understanding of complex evolutionary phenomena.

## Results

### Theory predicted SDC maintenance in strong segregation

To illustrate the basic concept of how self-destructive cooperation were maintained in strong segregation, we considered the most extreme case of cooperation, where all the benefits were utilized by the non-sacrificing cheaters in heterogenous groups composed of SDCs and cheaters. In this case, SDCs was assumed to entirely vanish 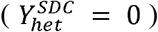 in these heterogenous subgroups, while the cheaters maintained 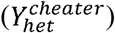, regardless of the initial ratio between SDCs and cheaters. According to traditional group selection theory, the maintenance of cooperators required positively correlated yield and initial ratio of SDC in heterogenous subgroups to balance the loss of SDC in every subgroup^11,12,27,28^. However, SDCs were assumed to vanish in all heterogeneous groups, independent of their initial ratio. This independence implied that there was no inter-group advantage to offset the extreme intra-group cost of the SDC sacrifice, making it impossible to maintain SDCs according to traditional group selection theory (see method and Extended Data Fig. 1a). In contrast, in homogenous subgroups of SDCs, individual SDCs benefited from the sacrifice of other SDCs, leading to a high population yield 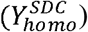. On the other hand, homogenous subgroup of cheaters vanished since there were no SDCs to protect the population under stress 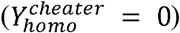 (Fig. 1a).

**Fig. 1.**
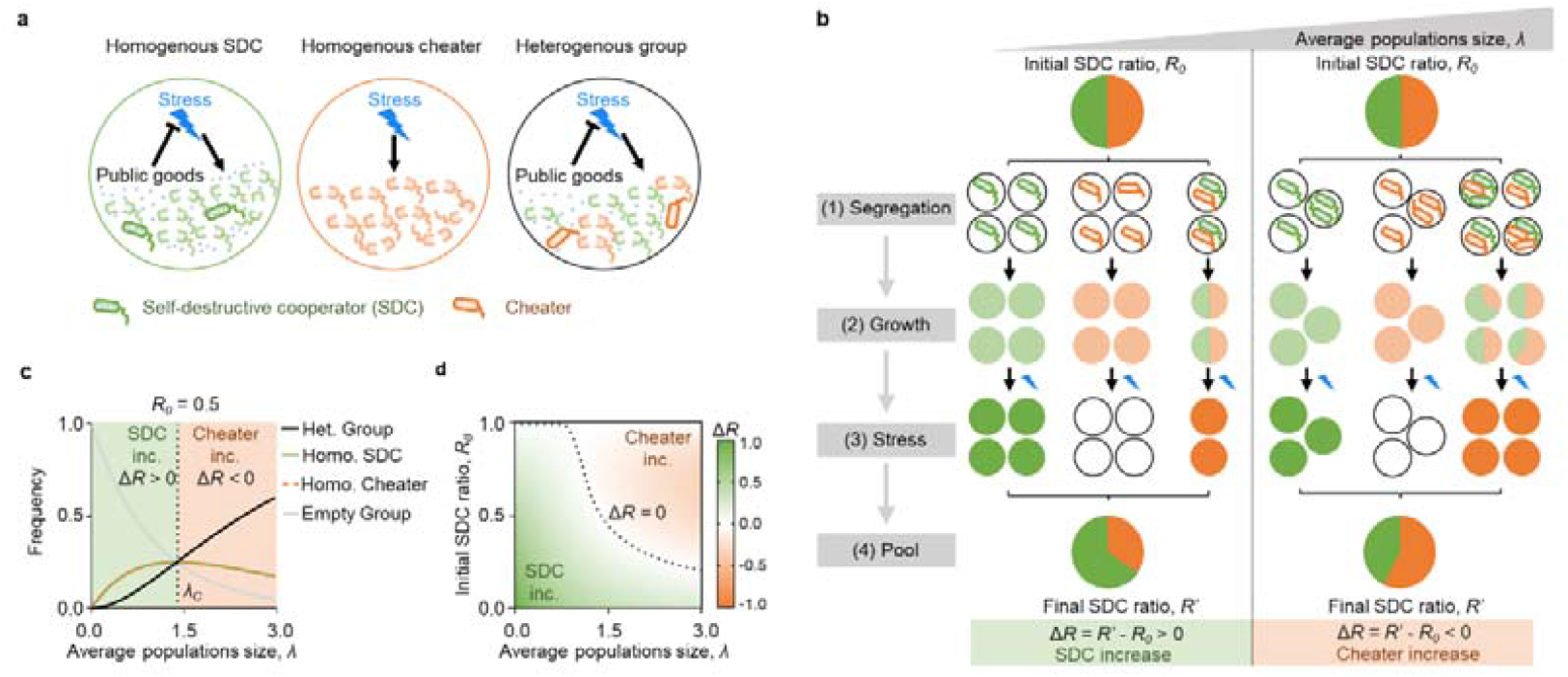
Theory predicted SDC maintenance under strong segregation. **a**, SDC (green strain) produced public goods (dots) that mitigated environmental stress (blue lightning bolts). Homogenous SDC groups (Homo. SDC) benefited from mutual sacrifice, leading to high yield after stress. Homogenous cheater groups (Homo. cheater) failed to survive without SDCs, while heterogenous groups allowed cheaters to exploit public goods, leading to cheaters proliferation. **b**, A mixed population of SDCs and cheaters was segregated into homogenous SDC, homogenous cheater, and heterogenous group under strong segregation. Smaller λ increased the *f*_*homo*_, boosting SDC survival and decreasing *f*_*het*_ that favored cheaters. Strong segregation thus facilitated an increase in Δ*R* (Δ*R* > 0) **c**, For *R*_*0*_ = 0.5, frequencies of homogenous and heterogenous groups were plotted against λ. With 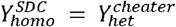, self-destructive cooperation maintenance required *f*_*het*_ ≤ 0.5 *f*_*homo*_ (dash line). **d**, The ΔR after the SGSP process, calculated using relation 2, was shown as a function of *λ* and *R*_*0*_. The dashed line represented the critical line Δ*R* = 0 separating SDC increase (SDC inc.) from cheater increase (Cheater inc.).

Here, the homogenous subgroups contributed to SDC offspring, while heterogenous subgroups contributed to cheater offspring. Then the maintenance of SDCs depended on the frequency of homogenous subgroups after segregation (*f*_*homo*_) as the frequency of heterogenous subgroups (*f*_*het*_) was dependent on *f*_*homo*_ (*f*_*het*_ = 1 - *f*_*homo*_) (Fig. 1b). Given the initial frequency of SDCs before segregation (*R*_*0*_), we could get the final ratio of SDCs after the SGSP procedures (*R*’) which could be expressed as:

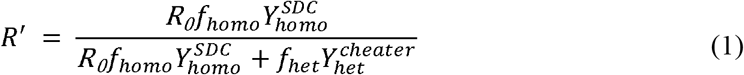

The condition for self-destructive cooperation maintenance was a positive change in SDC ratio (Δ*R* ≥ 0), with Δ*R* = *R*′ - *R*_0_. In a more simplified case where the yields of SDCs and cheaters were equal 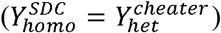, the condition of self-destructive cooperation maintenance became:

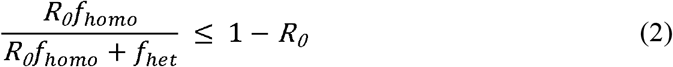

In the most simplified case of *R*_*0*_ = 0.5, this condition reduced to *f*_*het*_ ≤ 0.5 *f*_*homo*_.

In natural segregation cases, such as the division of biofilms, the formation of subgroups often followed the random sampling process, which led to a Poisson distribution of population sizes across the resulting subgroups^29,30^. Consequently, the proportions of homogenous and heterogenous subgroups were solely determined by the average population size *λ*, defined by the average cell number across all subpopulations. In this scenario, strong segregation favored the formation of homogenous SDC groups because smaller *λ*(stronger segregation strength) led to larger *f*_*homo*_ and smaller *f*_*het*_ thus facilitating an increase in the Δ*R* (Δ*R* ≥ 0) (Fig. 1c). Relation 2 also allowed the calculation of the critical segregation strength (*λ*_*c*_) required to maintain self-destructive cooperation by allowing 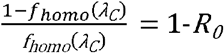. In the case of a balanced initial population (H_0_ _ 0.5), the critical segregation strength was approximately *λ*_*c*_ ≈ 1.5 (dashed line in Fig. 1c). Moreover, lower *R*_*0*_ necessitated a larger *λ*_*c*_ (weaker segregation strength) for the self-destructive cooperation to persist (Extended Data Fig. 2). This inverse relationship between the *R*_*0*_ and the *λ* on the Δ*R* was illustrated in a phase diagram (Fig. 1d).

Although our theoretical exploration showed that self-destructive cooperation could evolve in structured environments with strong segregation, experimental validation posed significant challenges. The primary challenge lay in creating a repeatable experimental framework that accurately modeled the SDC-cheater system. Additionally, population growth from 1 or 2 cells generated high variability in growth dynamics across a large number of subgroups. Therefore, a high-throughput pipeline with continuous monitoring and in-demand manipulation was required for the experiments^26^. In this paper, we overcame these challenges with synthetic bacterial strains and an automated biofoundary.

### Synthetic SDC-cheater system with engineered *E. coli*

To probe the evolution of self-destructive cooperation, we employed a biological synthetic system with two engineered strains of *E. coli* (Fig. 2a): one embodying the SDC phenotype and the other representing non-sacrificing cheaters. The SDC strain was engineered to undergo programmed cell death in the presence of antibiotics (6-APA). After cell death, the pre-expressed antibiotic-degrading enzyme (BlaM) was released into the environment, decreasing antibiotic concentration^2^. Meanwhile, the cheaters experienced cell lysis because of the antibiotic but did not provide any BlaM.

**Fig. 2.**
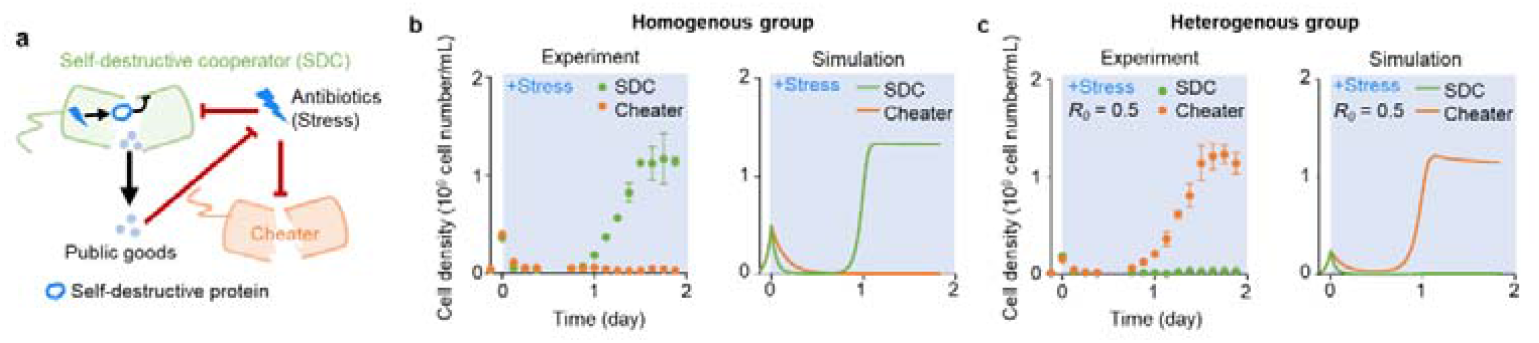
Engineered microbial SDC-cheater system. **a**, The SDC strain (green strain) underwent programmed cell death when exposed to antibiotic stress (blue lightning). This self-destruction released public goods (dots), such as the antibiotic-degrading enzyme BlaM, which mitigated the antibiotic stress. The cheater strain (red strain) underwent cell lysis without providing BlaM. **b-c**, Experiments and simulations of the growth dynamics of engineered SDCs and cheaters in the homogenous group (**b**) and heterogenous group with an initial SDC ratio of 0.5 (**c**), with 0.4 mg/ml 6-APA added at time 0. The error bar represented the standard deviation of three biological replicates. Data points exceeding three standard deviations above the mean were excluded from analysis.

In a homogenous group of SDCs, a minor subset inevitably survived environmental stress, facilitated by the public goods contributed by sacrificed SDCs, which served to mitigate the stress. Once the stress diminished to tolerable levels, the surviving SDCs regained their capacity to grow, ultimately giving rise to a new colony of SDCs with high yield 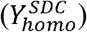. However, in a homogenous group of cheaters, the stress was not diminished effectively, resulting in a persistently inhibited cheater population at a reduced level yield 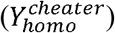 (Fig. 2b). In heterogenous groups, the imposition of external stress triggered the self-sacrifice of SDCs, which released public goods to mitigate the stress. Initially, the non-sacrificing cheaters were suppressed by the stress. However, they recovered and began to grow earlier than the surviving SDCs, allowing the cheaters to dominate the population 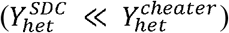 (Fig. 2c). The growth dynamics were then captured by a set of ordinary differential equations (ODE) (Extended Data Tab. 1 and methods). Furthermore, as long as the SDCs were able to generate sufficient public goods through self-sacrifice to lower the stress (*R*_*0*_ > 0.3) to a tolerable level, the yield of the cheaters became independent on *R*_0_ but only dependent on the nutrients in the environment (Extended Data Fig. 3). This phenomenon occurred because all SDCs were triggered to undergo self-sacrifice, leaving behind only negligible survivors that did not affect the yield of cheaters. In a typical antibiotic concentration of 0.4 mg/ml, we had 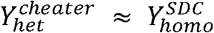 (Fig. 2), which allowed the application of relation 2.

### Validation of SDC evolution using automated biofoundry

To continuously monitor growth dynamics and introduce the antibiotic stress at a consistent time setting for subgroups with high growth viability, we employed an automated biofoundry to control the procedures precisely (Extended Data Fig. 4). This biofoundry utilized six devices, a robotic arm, two operating systems, and a flow cytometer, completing the SGSP process in 18 steps (see methods).

The *R*_0_ and *R*’ were manually measured by flow cytometry. During the automated high-throughput SGSP experiments, individual subgroups were pre-cultured in multiple 384-well plates to the stationary phase, and the cell density was monitored through a plate reader (Fig. 3a and Extended Data Fig. 5a-b). To ensure antibiotic stress was introduced uniformly at a similar cell density (0.1 < optical density (A600) < 0.3, Extended Data Fig. 5c), each subgroup was diluted to an A600 of approximately 0.04. After 75 min re-culturing, a certain concentration of the antibiotic was then added to all the subgroups (Extended Data Fig. 5d). The established automated workflow (Fig. 3b and methods) offered an 18-fold improvement over the manual operation, reducing the time required for experiments from 4 months to just one week. This efficiency was achieved by running three different sets of experiments simultaneously, enabling up to 18 automated workflows within a week while eliminating human errors and inconsistencies (see methods).

**Fig. 3.**
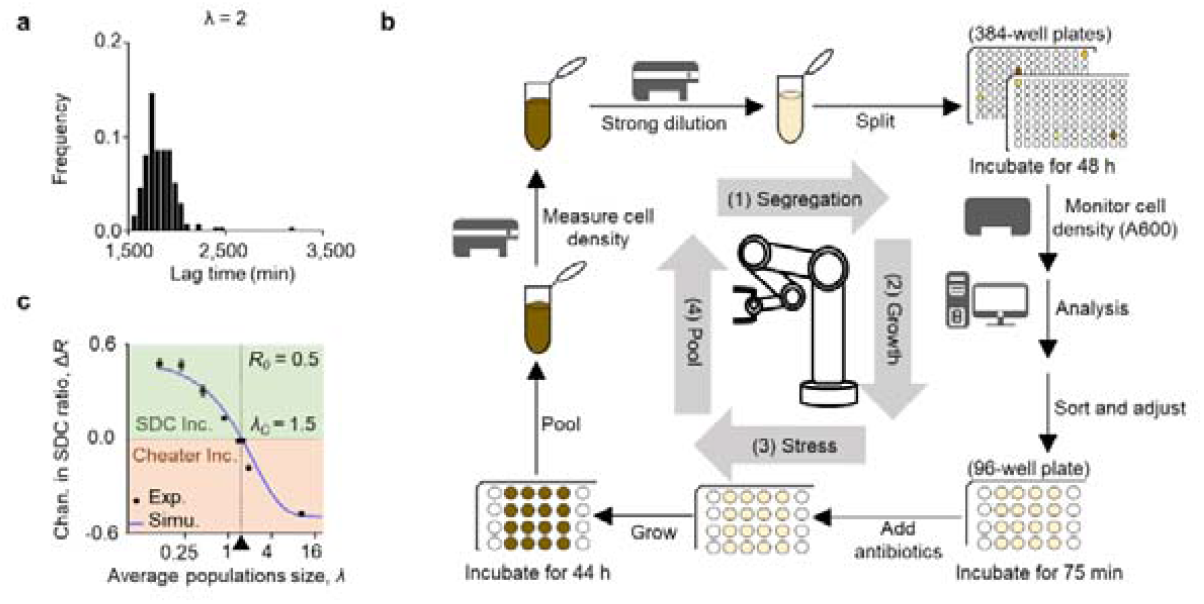
Biofoundry validation of SDC evolution under stronger segregation. **a**, Frequency distribution of the lag times of the population dynamics for 3∼4 colonies when λ = 2. The lag time measurement was detailed in the methods and Extended Data Fig. 5a. **b**, Illustration of an automated SGSP workflow with the biofoundry technology (see methods for details). **c**, Experimental and simulated results showed that the Δ*R* decreased with λ. A positive Δ*R* (Δ*R* > 0) indicated an increase in SDCs, while a negative Δ*R* (Δ*R* < 0) showed an increase in cheater. The error bar represented the standard deviation for three technical replicates, and the antibiotic stress was set at 0.4 mg/ml 6-APA. Exp., experiment; Simu., simulation.

Using this automated protocol and synthetic SDC-cheater system, we investigated the Δ*R* at various segregation strengths (characterized by *λ*). The findings revealed a distinct pattern: when the initial group contained 50% SDC individuals (*R*_*0*_ = 0.5), the *λ* threshold (*λc*) was 1.5, indicating Δ*R* > 0 when *λ* < 1.5 and *vice versa* (Fig. 3c and Extended Data Fig. 6). This aligned well with theoretical predictions, suggesting that the evolution of SDC depended on the relative frequencies of homogenous SDC and heterogenous groups (Extended Data Fig. 7a). Furthermore, we confirmed the ability of self-destructive cooperation to evolve and persist across multiple rounds of SGSP procedures (Extended Data Fig. 7b). Collectively, these results supported the theory that self-destructive cooperation could maintain and evolve in strong segregated environment.

### High stress facilitated SDC evolution

As the abovementioned theory, a simplified assumption was made: the yield of homogenous SDC groups was equal to that of heterogenous groups. However, this assumption did not hold when the stress level changed. We then investigated the impact of stress level (antibiotic concentration) on the yields of SDCs and cheaters. We found out that higher stress levels decreased cheaters’ yield within heterogenous groups (Fig. 4a). This change was attributed to the limited capacity of public goods produced by SDCs to degrade more antibiotics in these mixed populations. Conversely, homogenous SDC groups demonstrated a much higher ability to degrade antibiotics, rendering the yield of SDCs independent of the antibiotic concentration. This finding suggested that higher stress levels promoted the evolution of SDCs. Notably, the yield of SDCs in heterogenous groups and homogenous cheater groups was negligible across all antibiotic concentrations tested.

**Fig. 4.**
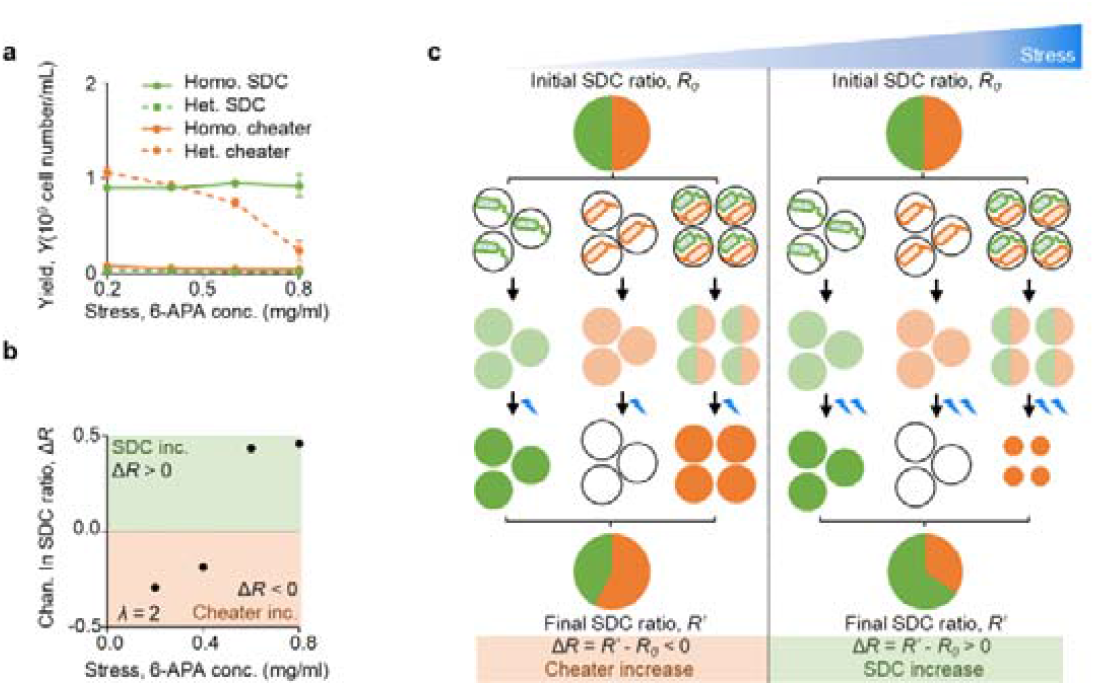
Stress facilitated SDC evolution in structured environments. **a**, Experimental data of the yields at varying stress levels (6-APA concentrations). The yield of SDC in the homogenous group (green solid line) remained constant, while the yield of the cheater in the homogenous group (red solid line) remained null. The yield of SDC in heterogenous groups (green dash line) remained null while the yield of cheater decreased with stress intensity (red dash line). Homo. SDC, homogenous SDC group; Het. SDC, heterogenous SDC group; Homo. cheater, homogenous cheater group; Het. cheater, heterogenous cheater group. **b**, The ΔR increased with stress intensity when *λ* = 2. ΔR ≥ 0 signified an increase in SDCs, whereas ΔR < 0 denoted an increase in cheaters. The error bar in panels a-b represented the standard deviation of three biological replicates. **c**, Illustration of how high stress facilitated SDC evolution. The reduced yield of heterogenous cheater groups under high stress created a selective pressure that favored the survival and proliferation of SDCs. Because SDCs produced public goods that alleviated stress, they were better equipped to survive and reproduce in stressful environments.

Further experiments employing the SGSP procedure with varying antibiotic concentrations revealed that even under weak segregation strength (*λ=* 2), higher antibiotic concentrations were required to promote SDC maintenance in structured environments (Fig. 4b). This phenomenon can be explained by the observation that higher antibiotic concentrations reduced the yield of cheaters, thereby diminishing their fitness advantage (Fig. 4c). The comprehensive model, which incorporated yield data under varying antibiotic concentrations (equation 1), allowed for theoretical recapitulation of the experimental findings. Notably, the effect of stress on SDC yields was directly implemented in the model without assuming any specific molecular mechanisms. We, therefore, concluded that high stress levels increased the advantage of homogenous SDC subgroups by decreasing the fitness of cheaters in heterogenous groups, thus creating a favorable environment for self-destructive cooperation to evolve.

## Discussion

This study shed light on the paradoxical emergence of self-destructive cooperation in structured environments. While self-destructive cooperation was considered an altruistic trait^2,31^, its evolutionary path remained puzzling. However, the discovery of similar gasdermin pathways in bacteria and animal cells suggested an evolutionary connection^32^, potentially supporting the “original sin” hypothesis of self-destructive cooperation origins, where death-related genes may have existed at the beginning of life^4^. Here, we proposed a mechanism where strong segregation and stressful environments promoted self-destructive cooperation evolution. Our theoretical deductions suggested that it occurred when the ratio of heterogenous SDC groups to homogenous groups fell below the initial cheater ratio (relation 2 and Fig. 1d). These theoretical predictions were validated using a synthetic SDC-cheater system in *E. coli*. Mimicking self-destructive cooperation and strong segregation through an automated biofoundry, we successfully proved self-destructive cooperation evolution in stressful environments.

Biofoundries represented a cutting-edge approach to laboratories, seamlessly integrating automation platforms for diverse workflows. Combining experimental approaches (mathematical modeling, computer-aided design, analytical software) with automated platforms enabled high-throughput workflows. This integration enhanced precision and stability through robotic arms and guide rails^35^. Our project utilized smaller cultivation volumes (microplates) to significantly increase throughput, addressing limitations of traditional methods and enabling rapid testing of numerous strains or variants. Although biofoundries have been applied in various fields, such as biosynthesis of plant-derived bioactive compounds^36^ and directed protein evolution^37^, their use in population genetics had been limited. This study pioneered the application of these technologies in population genetics, redesigning workflows for strong segregation conditions and demonstrating SDC evolution under such circumstances. The established automated workflow (Fig. 3b and methods) offered an 18-fold improvement over manual operations, enabling faster processing and eliminating human errors. This suggested a significant advancement of automated technology, highlighting its efficiency and reliability.

Our findings highlighted the dependence of self-destructive cooperation evolution on various environmental factors. Notably, the *λ* representing segregation strength, emerged as a critical factor in realistic scenarios with random sampling (Poisson distribution). Additionally, we observed a counterintuitive phenomenon: higher antibiotic concentrations, representing increased stress, facilitated SDC evolution. These insights were valuable for understanding the adaptive strategies employed by microorganisms in response to environmental stressors. Further exploration of SDC dynamics and its implications for microbial ecology held promise for advancing our understanding of cooperative evolution and designing intervention strategies to treat bacterial infections effectively.

In classical group selection theory, the Price equation was a pivotal analytical tool to evaluate the evolutionary potential of cooperative traits^10,33,34^. Under conventional segregation dynamics, wherein the population was uniformly allocated into subgroups with an equitably dispersed initial ratio of SDCs, the Price equation yielded a negative outcome, suggesting that SDCs could not be maintained through typical segregation mechanisms (Extended data Fig. 1a and method). Conversely, when strong segregation was imposed, homogenous groups dominated the genotypes of subgroups, which resulted in a pronounced positive covariance between fitness (*W*) and the initial SDC ratio (*cov* (*W,R*_*0*_)) and a low negative value of SDC fitness loss in each subgroup 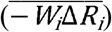. We recalculated the essential components of the Price equation as a function of the mean population size (*λ*), utilizing parameters established in Extended Data Fig. 1b-d. As *λ* increased (weaker segregation strength), *cov*(*W,R*_*0*_) decreased while 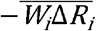 increased. The critical value where the two terms intersected was found at *λc* 1.5, which was consistent to our theoretical predictions and experimental observations (Fig. 1c & Fig. 3c). Thus, the persistence of SDCs under strong segregation did not violate the classic group selection theory, although this phenomenon had neither been demonstrated nor predicted before.

## Methods

### Conditions for self-destructive cooperation evolution

Extreme dilution, caused by random sampling of bacterial cells, could lead to strong segregation and a Poisson distribution of isolated groups with varying compositions ^29,30^. This scenario allowed calculation of the simplified frequency of both homogenous SDC groups: 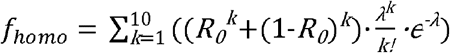 and heterogenous groups: 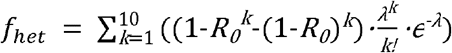 using equations that considered the *λ* and *R*_*0*_. Symbols were used to represent different parameters: 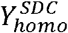 and 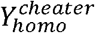 denoted the yield of homogenous SDC and cheater groups, respectively, while 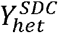 and 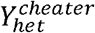 represented the yield of heterogenous groups containing SDCs and cheaters. Based on these parameters, a formula was derived to track the final SDC ratio over time: 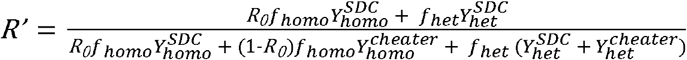. This formula was simplified because SDCs completely sacrificed themselves in heterogenous groups 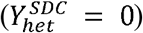. Additionally, the cheaters were all killed by the stress in homogenous cheater groups 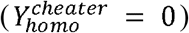. Furthermore, homogenous SDC or heterogenous cheater exhibited the same yield 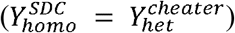 (Extended Data Fig. 3). With these simplifications, the formula became: 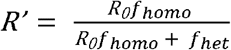, where *R*_*0*_*f*_*homo*_ and *f*_*het*_ represented the frequencies of homogenous SDC and heterogenous groups, respectively. The change in SDC ratio was represented by Δ*R* =*R*’ – *R*_*0*_. Thus, self-destructive cooperation evolution occurred when Δ*R* ≥ 0. Solving for this condition obtained Relation 2. The calculation for *f*_*homo*_ and *f*_*het*_ was truncated at *k* =10 as without affecting the accuracy of the outcome (Δ*R*) (Extended Data Fig. 8).

### ODE model for the synthetic SDC-cheater system

To capture the dynamic of the co-cultured strains, we modified the model first developed in Tanouchi 2012 MSB to characterize the growth curve of single strains^2^. Besides putting the equations of two strains together, we also added a nutrition equation to predict the maximal cell density that a system could reach, and also a resistance term of cell lysis to describe the resistant cells caused by phenotypic diversity. The full model could be written as follows:

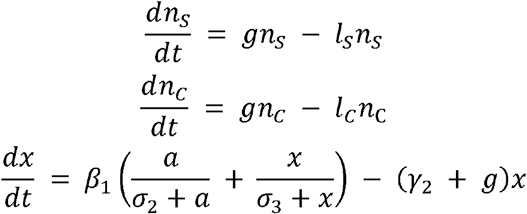

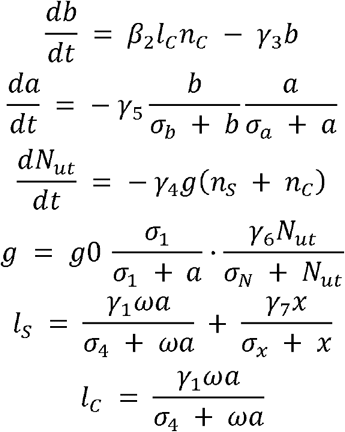

Where *n*_*S*_ *n*_*C*_ represented cell density of SDC and cheater; *x* was the intracellular concentration of E protein; *b* was the extracellular concentration of BlaM; *a* was the extracellular concentration of 6-APA; *N*_ut_ represented the nutrition left in the system; *g* was the growth rate of both cooperator and defector, as they were in a common growth condition, the growth rate was considered identical to both strain; *l*_*S*_ *l*_*C*_ were the lysis rate of SDC and cheater. The parameters used in this research were either measured in related references or chosen in a biochemical reasonable range, and they were listed and described in Extended Data Tab. 1. The initial condition was set to *n* (0) = 0.001,*x* (0) = 0, *b* (0) = 0 antibiotic 6-APA was added to the system at 6 h by letting a(0.4) = 0 → 1.

### Cell strains

SDC and cheater strains were derived from the *E. coli* SN0301 strain (*ampD1, ampA1, ampC8, pyrB, recA* and *rpsL*), which the *ampD1* mutation enabled hyper-induction of P_*ampC*_ in response to beta-lactam antibiotics^38^. The SDC strain was transformed with the pBlaM plasmid, pCSaE450C plasmid, and a GFP reporter. The cheater strain was transformed with pPROLar.A122 (Clonetech), pTS1^39^ and a mCherry reporter^2^. The pBlaM plasmid encoded the *blaM* gene under the control of the P_*lac/ara-1*_ promoter from pPROLar.A122, which had a p15A origin of replication. The pCSaE450C plasmid encoded the E gene under the P_*ampC*_ promoter through AmpR and was based on pTS1 with a pCDF13 origin of replication. The fluorescent reporters (GFP or mCherry) were carried on plasmids based on pZS31 with a pSC101 origin of replication.

### Culture Medium

Unless otherwise noted, Luria-Bertani (LB) medium and LBKM medium (10 g/L tryptone, 5 g/L yeast extract and 7 g/L KCl mediated by 100mM propanesulfonic acid and 5 M KOH at pH 7)^40^ were used for growth dynamic assays. Plasmids were maintained with 50 μg/ml spectinomycin, kanamycin and chloramphenicol. For every experiment, 100-fold 6-APA solution was prepared fresh by dissolving it in 1 M HCl. Appropriate concentrations of 6-APA and 1 mM IPTG were added to growth medium when applicable. All experiments were carried out at 37 □.

### Growth curve measurement

SDC and cheater strains were inoculated into LBKM medium for 18 ∼ 19 h (A600 was calibrated the same as ∼ 0.9). The overnight cultures were then mixed at various ratios to obtain the populations with *R*_*0*_ = 0 to 1. The mixed populations were further diluted 200-fold into 200 µL wells of LBKM medium (with 1 mM IPTG) on a 100-well honeycomb plate. For each condition, sextuple wells were implemented for the result reproduction. All the populations were then incubated for 4 ∼ 5 h with shaking until reaching A600 = 0.15 ∼ 0.2, after which an appropriate concentration of 6-APA was introduced into the corresponding wells to induce cooperation between the two strains. The wells without any 6-APA were considered as controls. Optical density measurements at a wavelength of 600 nm were performed using the Bioscreen C Automated Growth Curves Analysis System (FP-1100-C; Oy Growth Curves Ab) in a 100-well honeycomb plate. Data were collected at 15 min intervals for 48h.

### Characterization of the relative SDC ratio of the cells

At every given time point, a 10 µL aliquot of the cell population from three chosen replicate wells was diluted 100-fold with pre-chilled cell counting buffer (0.9 % NaCl with 0.12 % formaldehyde, filtered using a 0.22 µm filter). The diluted samples were then placed in an ice-water bath until cell counting. Before cell counting, the cell suspension was further diluted 10 times with 0.9 % NaCl (filtered using a 0.22 µm filter), for a total dilution factor of 10^−3^. Bacterial cell counting was performed with a flow cytometer (Beckman; CytoFLEX S), the flow rate and running time were 60 μL/min and 60 s, respectively. The cell density of SDC (with GFP) and cheater was obtained using a 488-nm laser and a 561-nm laser, respectively (Extended Data Fig. 9). The gain for FSC, SSC, FITC, and ECD channels was set to 500, 500, 1,500 and 1,500, respectively. At the final time point, the cell density, measured by either a plate reader or flow cytometer, was defined as the yield.

The SDC ratio was calculated by dividing obtained cell density of SDC to the total number of fluorescently labeled bacterial populations by flow cytometer. We defined the proportion of susceptible SDC before adding antibiotics as *R*_*0*_, and the proportion of SDC at the final time point as *R*’. The Δ*R* value was obtained by subtracting the proportion of SDC before adding antibiotics from the proportion of SDCs at the final time point.

### Characterization of the growth curve of the 384-well plate after strong dilution

SDC and cheater strains were inoculated into LBKM medium for 18 ∼ 19 h (A600 was calibrated the same as ∼ 0.9). The flow cytometer (Beckman; CytoFLEX S) was flushed with 0.9 % NaCl (filtered using a 0.22 µm filter) at a high speed (the flow rate was 240 μL/min) for more than 30 min to reduce the impact of blank events. Then the bacterial cell counting was performed with a flow cytometer using the same parameters as previously set (the flow rte and running time were 60 μL/min and 60 s, respectively). The bacteria were mixed in a 1:1 ratio directly on the flow cytometer. Following cell enumeration using the flow cytometer, the suspension was then strongly diluted with 0.9 % NaCl to achieve a *λ*value of approximately 2 in a 384-well plate. This dilution corresponds to an estimated concentration of 1,000 bacteria per 60 μL of suspension per well, based on post-dilution cell counts. 60 μL of the diluted bacterial suspension was added to 25 mL LBKM medium containing the corresponding antibiotics and dispensed into a 384-well plate. The plate was sealed with a film (breath-easy, Diversified Biotech), and A600 value were measured using a plate reader (Epoch 2; Biotek). Data were collected at 10 min intervals for 60 h.

### Statistical analysis of bacterial lag phase and yield

To understand how strong dilution impacts bacterial growth, we employed statistical analysis of the obtained growth curve data. For each well, we calculated the average A600 value between 300 and 1,000 min, representing a period of stable growth. This average served as a baseline to identify the end of the lag phase. The lag phase was considered to conclude when the A600 value surpassed this baseline by a factor of one (indicating significant growth) and exceeded a minimum threshold of 0.05. This point signified the transition to exponential growth. The lag time, the time required for bacteria to adapt to the lower cell density caused by dilution and initiate active growth, was then calculated as the duration from the start of the culture to the end of the lag phase.

In addition to the lag phase analysis, bacterial yield was evaluated based on the A600 value at the final time point.

### Biofoundry SGSP procedures for self-destructive cooperation evolution

Experiments were conducted using a robotic workcell housed within the Shenzhen Infrastructure for Synthetic Biology (SISB, Extended Data Fig. 4). This workcell contained a robotic arm on a 3.6-m track (Thermo Scientific, spinnaker mover), a liquid handling robot (Tecan, Freedom EVO 200) which was outfitted with a Roma robotic manipulation arm, a Liha 8-channel independent pipetter, a MCA96 with EVA adapter for 96-channel pipetting, a Te-Shake Silver shaker for high efficiency microplate oscillator. A microplate centrifuge (Agilent, G5582A), a microplate reader (Thermo Scientific, VarioskanLUX), an incubator (Thermo Scientific, Cytomat 2C 450 LIN Tos), a plate sealer (Kbiosystems, WASP), and a Xpeel seal pealer (Brooks Life Sciences, XP-A_230V) were used. Software for control and data integration across these devices was provided by Momentum (Thermo Scientific, Version 6.0.2). Fluent control software (Tecan, Version 2.7) managed the liquid handling robot functions, including pipetting programs, temperature control, and labware transportation. The robotic platform facilitated the evolution of self-destructive cooperation in recombinant *E. coli* through a series of automated steps: segregation, growth, stress, and pool. Here was an example of a biofoundry experiment setup for three different antibiotic treatments.

#### Segregation

1) Using flow cytometry, twice-activated overnight bacterial cultures were mixed in predefined ratios and diluted to specific concentrations with culture medium within a 1.5 mL Ep tube.
2) The diluted culture and a reservoir containing 150 mL of LBKM medium supplemented with antibiotics and IPTG were then placed in a tube carrier and Te-Shake in the Freedom EVO 200 liquid handler robot, respectively.
3) A precise volume was transferred from the Ep tube to the reservoir to achieve the desired final concentration. Before each pipetting step, the Te-Shake mixed the solution in the reservoir for 10 s and used a 96-channel pipetting to aspirate and dispense the bacteria solution 3 times to create a well-mixed bacterial solution.
4) Finally, the robot transferred the strong diluted bacteria solution into six designated 384-well plates.
5) Following the sealing, the plates were transferred to an incubator set at 37 °C and 1,000 rpm for incubation (supplemental video 1).

#### Growth

6) Following a 48 h incubation, a seal pealer removed the membranes from two out of the six 384-well plates (these two plates constituted one experimental set).
7) The Spark microplate reader then measured the yield (final cell density, A600) of the cultures, typically ranging from 0.4 to 0.8 (Extended Data Fig. 5a). Wells with A600 readings exceeding 0.4 were selected for further analysis.
8) A custom program calculated the volume of bacterial culture needed to dilute them to a common target density (*e*.*g*., A600 = 0.04) based on their A600 values at the stationary phase.
9) The Freedom EVO 200 liquid handler robot in SISB performed this dilution process. The first row of a 96-well plate served as a blank control.
10) Following dilution, the WASP plate sealer sealed the two 96-well plates, which were then transferred to an incubator set at 37 °C and 1,000 rpm for further culturing using the robotic arm.
11) The remaining four 384-well plates were processed in sets of two, following the same experimental procedures described above (supplemental video 2).

#### Stress

12) The bacterial cultures were incubated in 96-well plates at 37 °C and 1,000 rpm on Te-Shake Silver shakers for each set of two processed 384-well plates (refer to the previous section).
13) After reaching 75 min of incubation (Extended Data Fig. 5d), the cultures underwent stress treatment with antibiotics. Each set received a specific concentration of antibiotics.
14) Following the addition of the stress, the plates were sealed with a plate sealer and returned to the incubator for a further 44 h of incubation at the same temperature and shaking speed (supplemental video 3).

#### Pool

15) Following a 44 h incubation, the robotic arm retrieved each set of two processed 96-well plates (refer to the previous section).
16) A seal pealer removed the membranes from the plates.
17) 10 µL of bacterial culture in sets of two were collected from each well containing bacteria (excluding control wells) and pooled into a single Ep tube using the liquid handler (supplemental video 4).
18) This pooled culture was then analyzed using flow cytometry to determine cell number and ratio.

In parallel, the remaining bacterial solution was diluted back to the initial concentration used in the current experiment, preparing it for the next selection round.

Each round of the SGSP process, involving three different sets, took approximately 5 days. This setup allowed for up to three automated workflows to be completed in a single day, resulting in a total of nine different sets of experiments. By staggering the workflows across multiple days, a total of 18 workflows were conducted. This automated approach increased experimental efficiency 18-fold compared to manual methods, reducing the timeline from 4 months to just one week while eliminating human errors and inconsistencies.

### *λ* calculation methods

Two distinct methods were employed to determine the, and their performance was compared against an established simulation. In the first method (Method 1), bacterial cells were counted using flow cytometry, and *λ* calculated by averaging the number of bacteria per well based on the dilution factor. The second method involved counting empty wells in a 96-well plate and estimating *λ* using the Poisson distribution formula, which related the frequency of empty wells to the average population size (Extended Data Fig. 7a). Both methods were evaluated for accuracy and reliability, with results benchmarked against a simulation model to ensure robustness.

### Analyzing self-destructive cooperation evolution using the Price equation

The Price equation was expressed as: 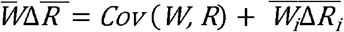. For self-destructive cooperation, we assumed that absolute fitness (*Wi*) was 0 for a homogenous cheater population. Conversely, other *R*_*0*_ received a fitness of 1. The *R’* was 1 for a homogenous group of cooperators and 0 for others. Notably, the Δ*R* decreased as the *R*_*0*_ varied, except for when it was 0 (Extended Data Fig.1 a-d). The Poisson distribution could lead to stochastic fluctuations of group compositions, resulting in more significant variances of *R*_*0*_ and *Wi*. Using parameters set previously, we calculated the *Cov* (*W, R*) and 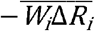. The point where these two values intersected was defined as *λ*_*C*_.

## Acknowledgments

We were grateful to the Shenzhen Infrastructure for Synthetic Biology for instrument support and technical assistance. The authors were grateful to Sihong Li, Pan Chu, Yi Zhang, Yue Yu, and Zhizhun Mo, as well as the other members of the Research Group, for helpful discussions on topics related to this work. This work was funded by the National Key Research and Development Program of China (2020YFA0908800, 2021YFA0910703), the Strategic Priority Research Program of the Chinese Academy of Sciences (XDB0480000), NSFC (32071417), Shenzhen Science and Technology Program (ZDSYS20220606100606013).

## Compliance with ethical standards

### Ethical approval

This article does not contain any studies with human participants or animals performed by any of the authors.

---

**Extended Data Fig. 1.**
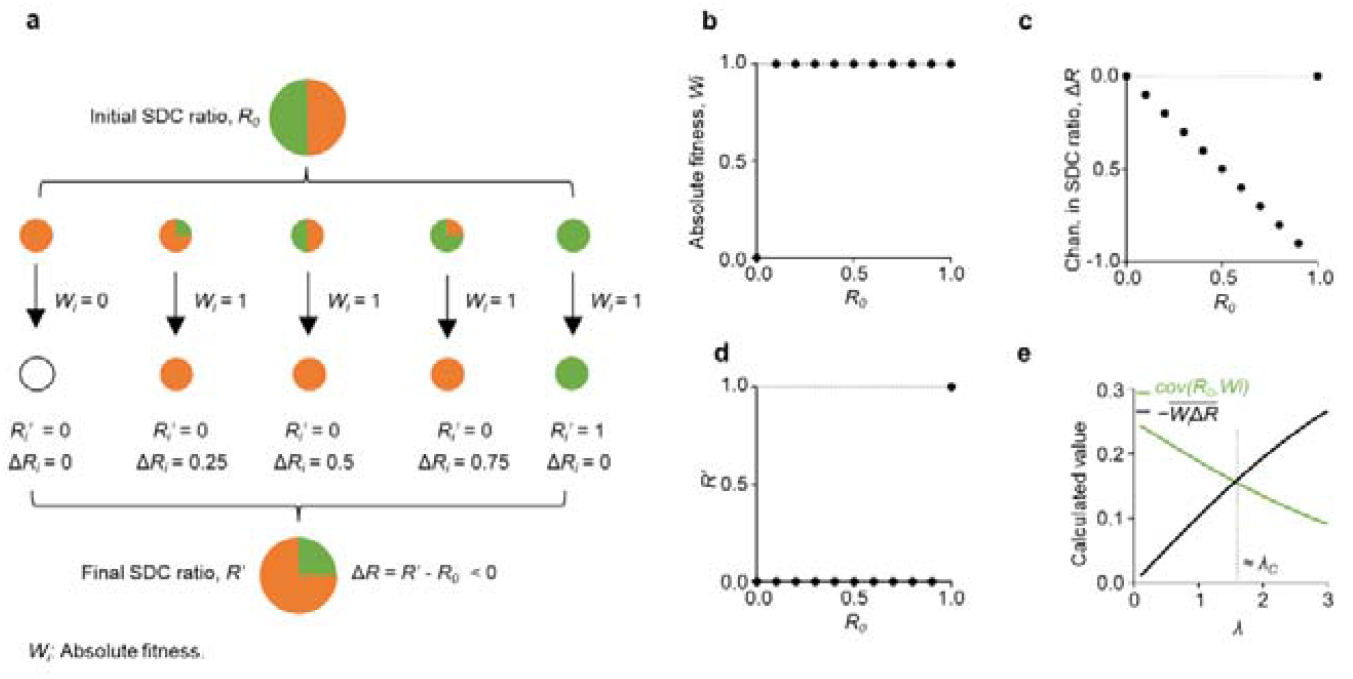
Traditional group selection theory failed to predict the evolution of SDC in structured environments. **a**, We employed the Price equation to analyze self-destructive cooperation evolution. Initially, each individual (*i*) carried a genotype (*R*_*i*_) designated as 1 for cooperators and 0 for cheaters (haploid). W_i_ varied based on the *R*_*0*_. The average genotype change 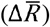 was expressed as: 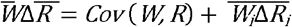. **b**, For SDC, *W*_*i*_ was 0 for a homogenous cheater, while it was 1 for other *R*_*0*_ values. **c**, The Δ*R* exhibited a decreasing trend with increasing *R*_*0*_, except for *R*_*0*_ = 1. **d**, The *R’* was 1 for a homogenous SDC and 0 for all other *R*_*0*_ values. Notably, even under the most extreme scenario where covariance between *R*_*0*_ and *W*_*i*_ equaled 1, the 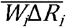 remained larger than covariance, hindering self-destructive cooperation evolution. **e**, To introduce significant variance, we proposed a mechanism involving extreme dilution of group populations (see methods). We calculated *Cov* (*W, R*) and 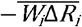 with decreasing *λ* (parameters set as described previously).

**Extended Data Fig. 2.**
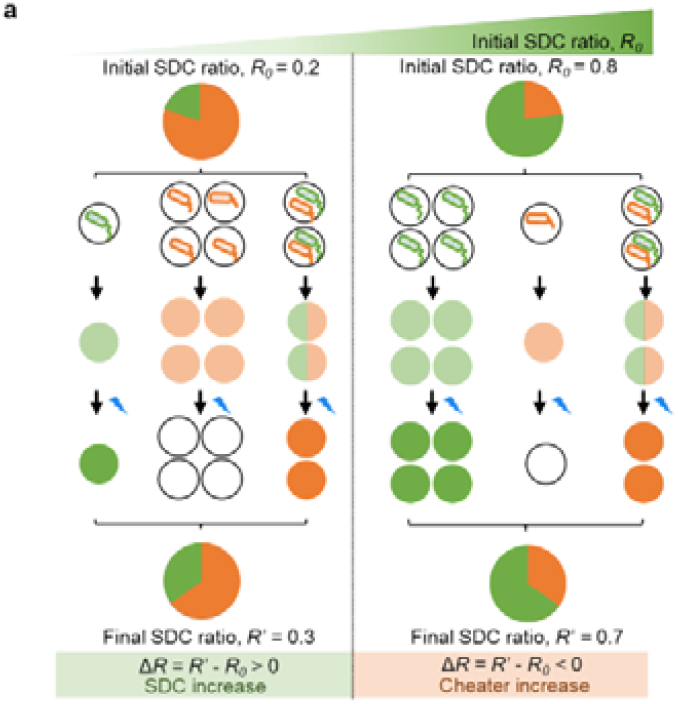
Lower *R*_*0*_ necessitated a larger *λ*_*C*_ for the SDCs to persist. In a fixed population structure, increasing the *R*_*0*_ increased the frequency of homogenous SDC groups while leaving the frequency of heterogenous groups unchanged. This implied that as *R*_*0*_ increases, the proportion of homogenous SDC groups relative to the total population also increased. However, when the groups were mixed after dispersion, this resulted in a lower *R’* compared to *R*_*0*_ for higher *R*_*0*_ values. Consequently, self-destructive cooperation could not evolve under these conditions.

**Extended Data Fig. 3.**
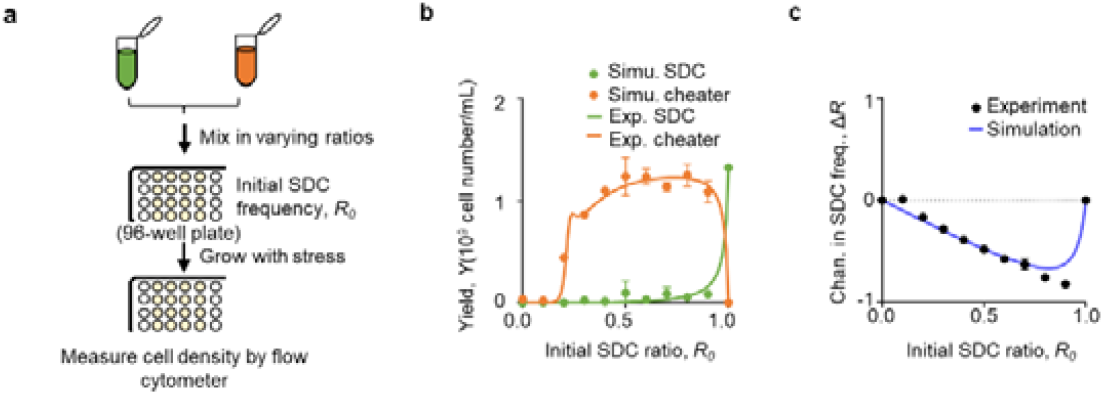
Validation of constructed microbial self-destructive cooperation system. **b**, This panel showed a schematic representation of the experimental setup for growing a mixture of SDC and cheater strains in varying ratios under stress conditions without segregation. **b**, This figure illustrated yield dynamics in different group compositions, comparing experimental results (points, n = 3) with ODE simulation results (lines) in the presence of 0.4 mg/ml 6-APA. **c**, This panel depicted the relationship between *R*_*0*_ and Δ*R* after stress induction. The result was consistent with the simulation outcome. Data represented the mean □ ± □s.d. for three biological replicates.

**Extended Data Fig. 4.**
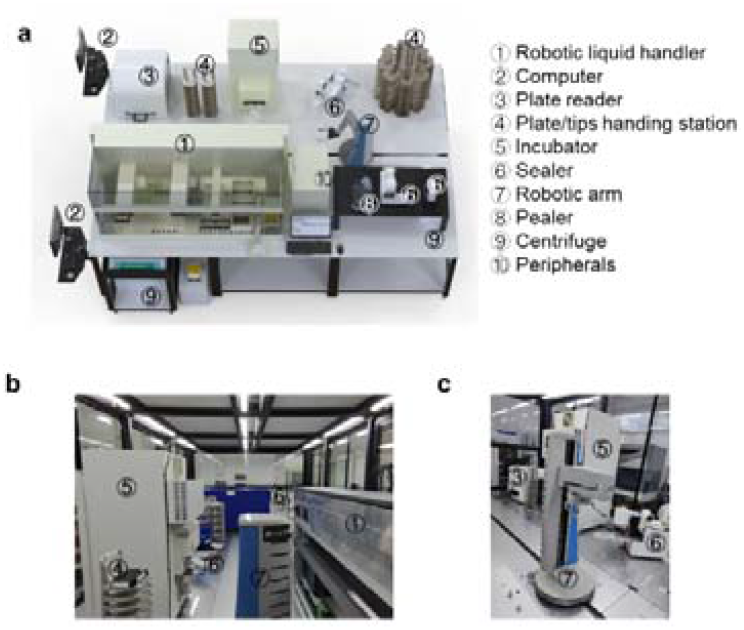
The Shenzhen Infrastructure for Synthetic Biology (SISB) biofoundry layout for SDC evolution. This figure illustrated the design and configuration of a biofoundry within the SISB specifically adapted for studying self-destructive cooperation evolution. **a**, The panel presented a schematic representation of the automated biofoundry. It depicted various core and peripheral instruments interconnected by a central robotic arm. A scheduling software managed the entire system, coordinating the operation of individual components. **b**-**c**, Panels b and c showcased physical drawings of the automation platform from two different perspectives.

**Extended Data Fig. 5.**
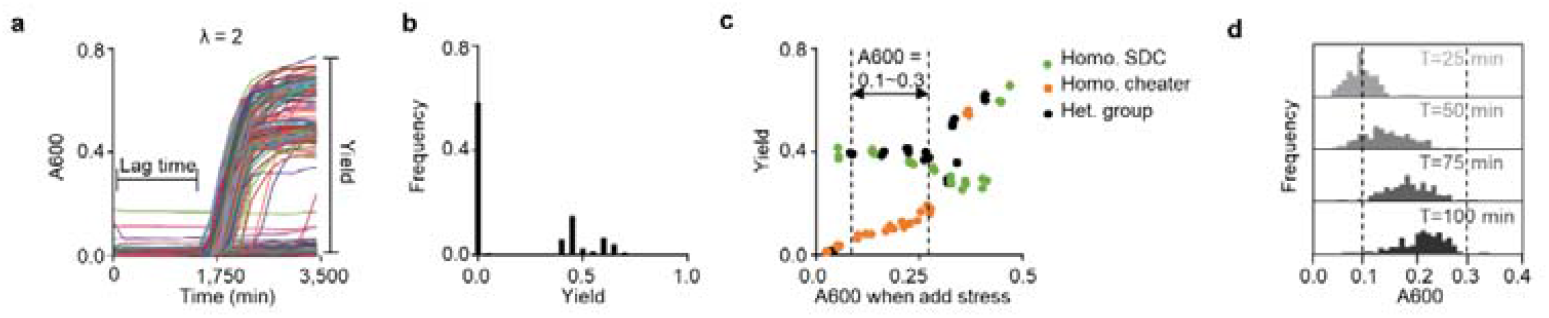
The necessity of an automated biofoundry for SGSP procedures. **a**, Variation in growth dynamics following dilution. Growth curves of multiple subpopulations when *λ* = 2 at 3,500 min (about 60 h) of culturing in a 384-well plate. Lag time was defined as the time to reach exponential growth (as shown in Fig. 3a), and yield was defined as the final cell density (A600). **b**, Frequency distribution histogram of the yield data when *λ* = 2, with intervals of 0.05 A600 units. **c**, This panel explored the impact of adding 0.4 mg/ml 6-APA at different A600 on yield. Two conditions were compared: monoculture (including homogenous SDC and homogenous cheater) and co-culture (heterogenous group, *R*_O_ = 0.5). The dashed line indicated that when stress was applied within this A600 range, the yields remained relatively stable. Homo. SDC, homogenous SDC group; Homo. cheater, homogenous cheater group; Het. group, heterogenous group. **d**, After the growth step, bacteria were re-cultured in a shaker, and their cell density was measured using a microplate reader within the automated biofoundry every 25 min. The dashed line indicated that the A600 was within the target range for adding the antibiotic.

**Extended Data Fig. 6.**
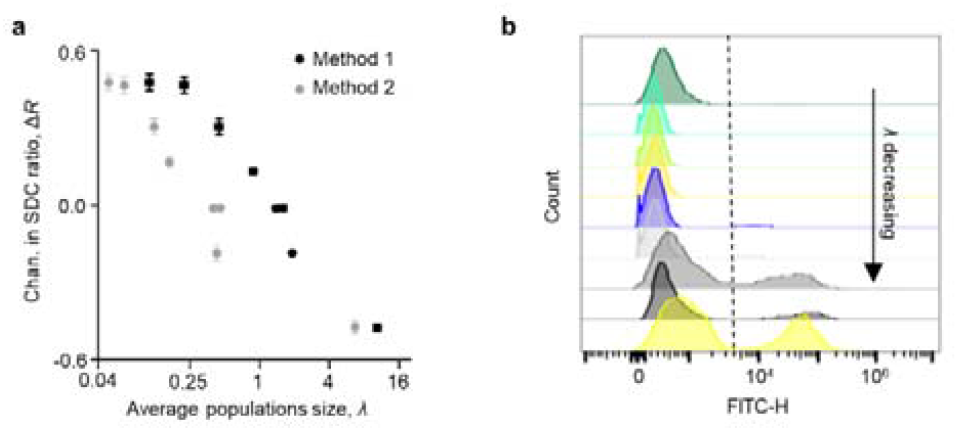
calculation methods and final SDC ratio distribution. **a**, Two methods were used to determine the *λ* when *R*_O_ = 0.5 (see methods). Method 1, indicated by the black dots, was believed to be more accurate due to the potential for underestimation in Method 2 (grey dots). This underestimation was likely caused by strong dilution impacts during the experiment, which may have hindered the recovery of some bacteria. Data represented the mean □ ± □s.d. for three technical replicates. **b**, Distribution of *R’* across varying *λ* values using flow cytometer.

**Extended Data Fig. 7.**
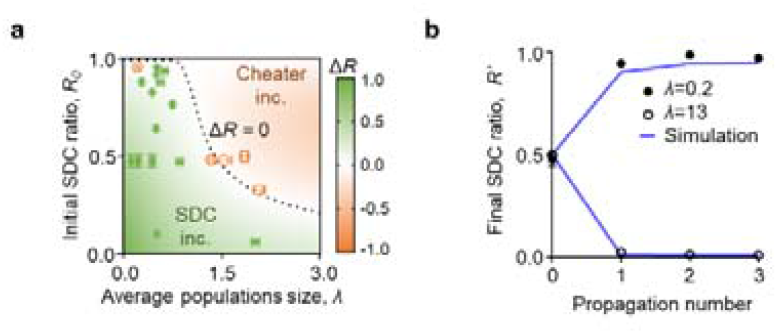
Confirmation of SDC evolution in small population sizes. **a**, Validation of the Δ*R* after the SGSP process, calculated according to equation 1, as a function of *λ* and *R*_*0*_ (as shown Fig. 1d). **b**, Comparison of experimental results (points, n = 3) with simulation results (lines) calculated using relation 2, across multiple cultivation rounds in either strong segregation (*λ* = 0.2) or weak segregation (*λ* = 13), in the presence of 0.4 mg/ml 6-APA with *R*_0_ = 0.5. Data represented the mean □± □s.d. for three technical replicates.

**Extended Data Fig. 8.**
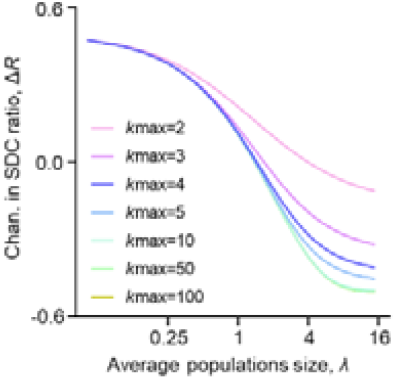
Truncated relation 2 model the summation at a specific *k* value. This involved truncating the frequency calculation formulas for *f*_*homo*_ and *f*_*het*_ group. The range of *k*max was examined from 2 to 100. Traditionally, the equation considered an infinite number of possibilities. Here, the summation was stopped at a specific point (*k*max) without affecting the accuracy of the outcome (Δ*R*). Thus, the *k* value in Relation 2 could be truncated at 10.

**Extended Data Fig. 9.**
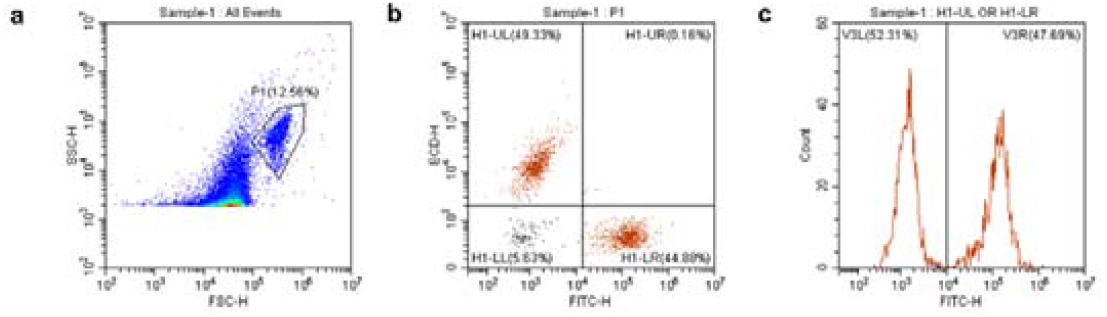
Gating strategy for flow cytometry analysis. **a**, Particles within region P1 were identified as bacterial cells based on forward scatter height (FSC-H) and side scatter height (SSC-H) characteristics. The established *E. coli* self-destructive cooperation system was labeled with a constitutively expressed green fluorescent protein (GFP) and red fluorescent protein (mCherry), respectively. This allowed quantification of their ratio through flow cytometry analysis. An example flow cytometry plot with a mixed population was shown. **b**, The plot of FITC-H versus ECD-H further differentiated the populations based on their respective fluorescent signals. Cells within H1-UL and H1-LR sub-populations were designated as cheater and SDC strains, respectively. H1-UL represented the GFP-negative, mCherry-positive population (cheaters), while H1-LR represented the GFP-positive, mCherry-negative population (SDCs) based on their fluorescence profiles. **c**, By comparing the number of SDC cells to the total number of fluorescently labeled bacterial populations, the *R*_*0*_ was obtained.

**Extended Data Tab. 1.** Model parameters of ordinary differential equations Parameter Symbol Value Reference.

| Parameter | Symbol | Value | Reference |
| --- | --- | --- | --- |
| Growth rate | $g_0$ | 0.9 /h | Calculated by experiment |
| Half-maximal constant for a degradation of 6-APA by beta-lactamase | $\sigma_b$ | 0.28 molecules/mL | |
| Half-maximal constant for a degradation by 6-APA | $\sigma_a$ | 0.1 mM | |
| E protein synthesis rate | $\beta_1$ | 18 molecules/cell | Copy number of pColE1 strain |
| BlaM production rate | $\beta_2$ | 10 molecules/cell | Copy number of p15A strain |
| Maximum cell lysis rate of cheater | $\gamma_1$ | 0.48 /h | Calculated by experiment |
| Degradation rate of E protein | $\gamma_2$ | 200 molecules/h | |
| Intrinsic degradation rate of BlaM | $\gamma_3$ | 0.09 molecules/h | |
| Yield nutrition consumption | $\gamma_4$ | 0.8 OD/mM | |
| Max degradation rate of 6-APA by BlaM | $\gamma_5$ | 0.06 mM/h | |
| Max nutrition -dependent growth rate | $\gamma_6$ | 1.5 mM/h | |
| Maximum cell lysis rate of SDC | $\gamma_7$ | 2.6 /h | Calculated by experiment |
| Half-maximal constant for growth inhibition of 6-APA | $\sigma_1$ | 0.02 mg/mL | |
| Half-maximal constant for E protein synthesis by 6-APA | $\sigma_2$ | 32.6 | Relative BlaM expression of $P_{ampC}$ in an $ampD^+$ strain (SN0301) to SN03 was 65.2<sup>38</sup>. |
| Half-maximal constant for E protein positive feedback | $\sigma_3$ | 2 | |
| Half-maximal constant for 6-APA-dependent lysis | $\sigma_4$ | 2 | |
| Half-maximal constant for nutrition-dependent growth | $\sigma_N$ | 0.6 mM | Max nutrition was 1.2 mM |
| Half-maximal constant for BlaM - dependent lysis | $\sigma_x$ | 0.004 molecules/cell | |
| Effective killing efficiency of antibiotic | $\omega$ | 10 | |

## Reference

1 Ackermann, M., Stecher, B., Freed, N. E., Songhet, P., & Doebeli, M. Self-destructive cooperation mediated by phenotypic noise. Nature 454, 987–990 (2008).

2 Tanouchi, Y., Pai, A., Buchler, N. E. & You, L. Programming stress-induced altruistic death in engineered bacteria. Mol. Syst. Biol. 8, 626 (2012).

3 Lee, H. H., Molla, M. N., Cantor, C. R. & Collins, J. J. Bacterial charity work leads to population-wide resistance. Nature 467, 82–85 (2010).

4 Ameisen, J. C. On the origin, evolution, and nature of programmed cell death: a timeline of four billion years. Cell Death Differ. 9, 367–393 (2002).

5 Bell, G. Selection : the mechanism of evolution. 367–368 (Oxford Univ. Press, 2008).

6 Radosevich, J. Apoptosis and Beyond : the many ways cells die. 237–288 (Wiley, 2018).

7 Cascales, E. et al. Colicin biology. Microbiol. Mol. Biol. Rev. 71, 158–229 (2007).

8 Nouvian, M., Reinhard, J. & Giurfa, M. The defensive response of the honeybee Apis mellifera. J. Exp. Biol. 219, 3505–3517 (2016).

9 Singer, M. et al. The third international consensus definitions for sepsis and septic shock (Sepsis-3). JAMA 315, 801–810 (2016).

10 Price, G. R. Selection and Covariance. Nature 227, 520–521 (1970).

11 Chuang, J. S., Rivoire, O. & Leibler, S. Simpson’s paradox in a synthetic microbial system. Science 323, 272–275 (2009).

12 Chuang., J. S., Rivoire, O. & Leibler, S. Cooperation and Hamilton’s rule in a simple synthetic microbial system. Mol. Syst. Biol. 6, 398 (2010).

13 Wilson, D. S. A theory of group selection. Proc. Natl Acad. Sci. USA 72, 143–146 (1975).

14 Nedelcu, A. M., Driscoll, W. W., Durand, P. M., Herron, M. D. & Rashidi, A. On the paradigm of altruistic suicide in the unicellular world. Evolution 65, 3–20 (2011).

15 Durand, P. M., Sym, S. & Michod, R. E. Programmed cell death and complexity in microbial systems. Curr. Biol. 26, R587–R593 (2016).

16 remer, J. et al. Cooperation in microbial populations: theory and experimental model systems. J. Mol. Biol. (2019).

17 Nadell, C. D., Drescher, K. & Foster, K. R. Spatial structure, cooperation and competition in biofilms. Nat. Rev. Microbiol. 14, 589–600 (2016).

18 Harcombe, W. Novel cooperation experimentally evolved between species. Evolution 64, 2166–2172 (2010).

19 Dobay, A., Bagheri, H. C., Messina, A., Kümmerli, R. & Rankin, D. J. Interaction effects of cell diffusion, cell density and public goods properties on the evolution of cooperation in digital microbes. J. Evol. Biol. 27, 1869–1877 (2014).

20 MacLean, R. C. & Gudelj, I. Resource competition and social conflict in experimental populations of yeast. Nature 441, 498–501 (2006).

21 Pande, S. et al. Privatization of cooperative benefits stabilizes mutualistic cross-feeding interactions in spatially structured environments. ISME J. 10, 1413–1423 (2016).

22 Kreft, J. U. Biofilms promote altruism. Microbiology (Reading) 150, 2751–2760 (2004).

23 Melbinger, A., Cremer, J. & Frey, E. The emergence of cooperation from a single mutant during microbial life cycles. J. R. Soc. Interface. 12, 20150171 (2015).

24 Rumbaugh, K. P. & Sauer, K. Biofilm dispersion. Nat. rev. Microbiol. 18, 571–586 (2020).

25 Granato, E. T., Meiller-Legrand, T. A. & Foster, K. R. The evolution and ecology of bacterial warfare. Curr. Biol. 29, R521–R537 (2019).

26 Alexander, H. K. & MacLean, R. C. Stochastic bacterial population dynamics restrict the establishment of antibiotic resistance from single cells. Proc. Natl Acad. Sci. USA 117, 19455–19464 (2020).

27 Xavier, J. B. Social interaction in synthetic and natural microbial communities. Mol. Syst. Biol. 7, 483 (2011).

28 Mysterud, I. Unto others: The evolution and psychology of unselfish behavior. Popul. Environ. 21, 581–588 (1999).

29 Hamilton, W. D. The evolution of altruistic behavior. Am. Nat. 97, 354–356 (1963).

30 Haldane, J. B. The causes of evolution. Vol. 5 (Princeton Univ. Press, 1990).

31 Tanouchi, Y., Lee, A. J., Meredith, H. & You, L. Programmed cell death in bacteria and implications for antibiotic therapy. Trends Microbiol. 21, 265–270 (2013).

32 Johnson, A. G. et al. Bacterial gasdermins reveal an ancient mechanism of cell death. Science 375, 221–225 (2022).

33 Page, K. M. & Nowak, M. A. Unifying evolutionary dynamics. J. Theor. Biol. 219, 93–98 (2002).

34 Damore, J. A. & Gore, J. Understanding microbial cooperation. J. Theor. Biol. 299, 31–41 (2012).

35 Wei, Y., Ji, X., & Cao, M. Engineering biology for microbial biosynthesis of plant-derived bioactive compounds 257–283 (Acad. Press, 2024).

36 Yuan, Y. et al. Efficient exploration of terpenoid biosynthetic gene clusters in filamentous fungi. Nat. Catal. 5, 277–287 (2022).

37 Yu, T., Boob, A. G., Singh, N., Su, Y. & Zhao, H. In vitro continuous protein evolution empowered by machine learning and automation. Cell Syst. 14, 633–644 (2023).

38 Lindberg, F., Lindquist, S. & Normark, S. Inactivation of the ampD gene causes semiconstitutive overproduction of the inducible Citrobacter freundii beta-lactamase. J. Bacteriol. 169, 1923–1928 (1987).

39 Sohka, T. et al. An externally tunable bacterial band-pass filter. Proc. Natl Acad. Sci. USA 106, 10135–10140 (2009).

40 Balagadde, F. K., You, L., Hansen, C. L., Arnold, F. H. & Quake, S. R. Longterm monitoring of bacteria undergoing programmed population control in a microchemostat. Science 309, 137–140 (2005).

